# A cocktail of rapamycin, acarbose and phenylbutyrate prevents age-related cognitive decline in mice by altering aging pathways

**DOI:** 10.1101/2022.09.07.506968

**Authors:** Zhou Jiang, Qianpei He, Warren Ladiges

## Abstract

Aging is a primary risk factor for cognitive dysfunction and exacerbates multiple biological processes in the brain, including but not limited to nutrient sensing dysregulation, insulin sensing dysfunction and histone deacetylation. Therefore, pharmaceutical intervention of aging targeting several distinct but overlapping pathways provides a basis for testing combinations of drugs as a cocktail. A recent study showed that middle-aged mice treated with a drug cocktail of rapamycin, acarbose, and phenylbutyrate for three months had increased resilience to age related cognitive decline. This finding provided the rationale to investigate the comprehensive transcriptomic and molecular changes within the brain of mice that received this cocktail treatment or control substance. Transcriptome profiles were generated through RNA sequencing and pathway analysis was performed by gene set enrichment analysis to evaluate the overall RNA message effect of the drug cocktail. Molecular endpoints representing aging pathways were measured through immunohistochemistry to further validate the attenuation of brain aging in the hippocampus of mice that received the cocktail treatment, each individual drug or controls. Results indicated that biological processes that enhance aging were suppressed, while autophagy was increased in the brains of mice given the drug cocktail. The molecular endpoint assessments indicated that treatment with the drug cocktail was overall more effective than any of the individual drugs for relieving cognitive impairment by targeting multiple aging pathways.

## Introduction

Aging is a complex process involving the change of multifactorial pathways in all organs of the body including the brain. Aberrant progression of several critical pathways accelerates chronological aging. Pathways including mammalian target of rapamycin (mTOR) signaling, insulin signaling and histone deacetylation [1] are respectively and individually targeted by rapamycin (Rap), acarbose (Acb) and phenylbutyrate (Pba), with each drug individually shown to have anti-aging effects in mice. A combination of these three drugs as a cocktail might be expected to result in a more robust outcome on preventing or slowing aging in the brain due to targeting multiple aging pathways, which each individual drug is not able to do [2].

Rap is an antibiotic clinically approved for treating organ transplant patients and neoplastic conditions [3,4]. It has a high oral bioavailability and readily crosses the blood brain barrier. Rap blocks mTOR1, a protein shown to integrate signals from growth factors and nutrients to control protein synthesis [5]. Mechanistically, Rap downregulates mTOR1 signaling, leading to a phenocopy of caloric restriction [6]. A previous National Insititute on Aging Intervention Testing Program (ITP) study demonstrated that Rap feeding increased lifespan even when started late in life [7]. Acb is a common type 2 diabetes medication used for glucoregulatory control by delaying the digestion of complex carbohydrates. Specifically, Acb reversibly inhibits α-glucosidases within the intestinal brush border and promotes postprandial blood glucose regulation [8]. A previous ITP study showed long term treatment of Acb in heterogeneous background mice extended both medium and maximum lifespan [9]. Pba is clinically approved as an ammonia scavenger for urea cycle disorders in children. Pba is a broad inhibitor to class I and II histone deacetylates which enhance histone acetylation and protein folding as well as decreases endoplasmic reticulum stress [10]. Extensive studies demonstrated that increases in histone acetylation globally elevate transcription and can be beneficial at old age as it contributes to reversion of age-dependent decline in stress response, DNA reparation and other genes involved in maintenance of homeostasis [11].

Middle-aged mice treated for three months with a drug cocktail consisting of Rap, Acb and Pba showed improvements in multiple aspects of health span [12] including significant delays in the appearance of cognitive impairment. Learning parameters were measured by a spatial navigation learning task [13]. Old mice that received cocktail and Rap showed enhanced learning abilities compared to all other cohorts. Rotarod, which exploits a mouse’s natural avoidance to height, was used to assess motor coordination [14]. Only mice treated with the cocktail showed significantly better performance. Both tests indicated that the drug cocktail was phenotypically effective suggesting a more robust effect than any individual drug.With the phenotypic observation on improved cognitive function, we posited that the drug cocktail was able to retard brain aging by targeting multiple biological processes, but the exact processes and pathways were not known. To investigate the pathway and molecular targets of aging in the brain from mice treated with drug cocktail or control, multivariate discriminant analysis was conducted for this study.

In order to identify age associated pathways altered by the cocktail in the brain, we explored transcriptomic profiles through RNA sequencing (RNA-seq). Pathways reported to be targeted by the drugs in the cocktail are mTOR1 signaling, insulin signaling and histone deacetylase binding, which represent three major pillars that are aggravated with increasing age including nutrient sensing, insulin signaling intensity, and epigenetic alternations. In addition, it was of interest to see if additional pathways of aging including DNA damage, inflammation, senescence, and autophagy [1] might also be alleviated by the cocktail.

Validation of these transcriptomic endpoints was done to demonstrate the expression levels of corresponding proteins using Immunohistochemistry (IHC). Digital image quantification was utilized to evaluate the intensity level from staining.

## Methods

### Mouse brain tissue

All tissues used for this study were obtained from archived brain samples of an NIA funded preclinical drug study, which compared aging phenotypes between the drug cocktail diet treated C57BL/6 mice with mice receiving individual drug diet or control diet. In this study, mice that received cocktail diet, each individual drug or control chow were tested.Mice were euthanized via carbon dioxide asphyxiation followed by decapitation. All brains were harvested and collected within 2 mins after decapitation. Brains were sectioned sagittally through midline. The parietal-temporal lobe from left sagittal hemisphere was extracted into a sterilized vial and snap frozen into liquid nitrogen before transferred to -80oC freezer for long term storage. The right sagittal hemisphere was fixed in 10% neutral buffered formalin (NBF) for 72 hours then embedded in paraffin wax through the University of Washington Department of Comparative Medicine’s Histology Lab.

### RNA-seq

Total RNA was extracted from homogenized parietal-temporal lobe tissue using Qiagen RNeasy kits (Cat. No.74106). RNA purity was measured by Nanodrop spectrophotometer and RIN value obtained through 2100 Bioanalyzer. Library construction and sequencing were done through Quick Biology Inc., using the high throughout genome sequencing platform NovaSeq 6000 with 15M paired reads. In brief, the raw RNA-seq data went through FASTQ quality control and clean up and was aligned to UCSD C57BL/6 mice RNA data bank for further analysis.

Quantitative reverse transcription PCR (q-PCR). RNA preparation for q-PCR was performed in the same manner as for RNA-seq. cDNA samples were generated by OligodT (High-capacity cDNA Reverse Transcription Kit; Thermo-Fisher Cat. 4368813) reverse transcription using 5ng RNA following the manufactural standard protocol. qPCR was carried out using SYBR Green Master Mix (PowerTrack SYBR Green Master Mix, Thermo-Fisher Cat. A46109) and primer pairs. β-actine was selected as a house keeping gene for quantification and all q-PCR results were analyzed through ΔΔCt method. Primer sequences are listed as follows: Cdkn2a (p16Ink4a) Fwd 5′-CCCAACGCCCCGAACT-3′, Cdkn2a (p16Ink4a) Rev 5′-GCAGAAGAGCTGCTACGTGAA-3′; β-actine Fwd 5′-CACCATTGGCAATGAGCGGTTC-3′, β-actine Rev 5′-AGGTCTTTGCGGATGTCCACGT -3′.

Gene set enrichment analysis (GSEA). GSEA is a useful tool to interpretate high throughout expression studies and identify insights into biological pathways underlying a given phenotype [15]. The DESeq R package was used for preparing differential expression transcriptome data across all sequencing cohorts. The GSEA algorithm scores a gene set according to how the genes in it represent increases or decreases on average in response to the regulation strength. The groups of pathways selected for GSEA analysis were predefined by previous studies [1] as major pillars of mammalian aging. Reference gene sets were extracted through GO or KEGG and GOMF database with strong validation from previous literature, and a filter was set between 15 to 500 genes. The chip platform applied to GSEA was mice gene symbols remapped to human gene orthologues for better translational results. The reference knowledge-based gene sets were extracted from gene ontology and KEGG terms containing gene identifiers and corresponding expression values. The GSEA analysis was done using GSEA software version 4.2.1 and the database was derived from the Molecular Signatures Database (MSigDB v5.0). GSEA was based on K¬-S test which weighs each observed gene within the gene set based on a given cumulative density function.

### Immunohistochemistry (IHC)

Brain tissue of 6 mice was randomly selected from each group. Brain from 8-month-old mice was measured for each tested endpoint as a negative control reference. 4μm formalin fixed paraffin embedded unstained slides were used for staining. Slides were rehydrated with xylene and a series of decreasing concentration of ethanol in water solution. Slides then went through 20mins heat medicated antigen retrieval (98 oC) in 1X citric acid buffer solution (PH 6.0). Slides applied with P21 as primary antibody were immersed in EDTA buffer (PH 8.0) for antigen retrieval. Sections then blocked with endogenous peroxidase plus 3% H2O2, serum, avidin and biotin blocking reagents (HRP-DAB Cell &Tissue Staining Kit, R&D Systems) and washing in TBST in between each blocking step. Sections were then incubated in primary antibodies diluted in TBST solution overnight in a 4 oC humidified chamber. The dilution concentration for each antibody are as follows: HDAC2 1/500 (Abcam, Cat. ab7029), MCP1 1/200 (Novus Biologicals, Cat. NBP1-07035), TNFα 1/400 (Abcam, Cat. ab1793), ATG5 1/500 (Novus Biologicals, Cat. NB110-53818), IL6 1/400 (Abcam, Cat. ab208113), P21 1/50(Abcam, Cat. ab188224), 1/500 γH2AX (Abcam, Cat. ab26350). Slides were rinsed thoroughly before incubating in biotinylated secondary antibody for 30mins at room temperature. After the washing step, sections were incubated with conjugated high sensitivity streptavidin house radish peroxide (HSS-HRP) for 30mins followed by washing steps. DAB chromogen was applied to slides for 20min until desired intensity of staining was reached. Slides were then dehydrated and mounted for imaging and analysis.

### QuPath analysis

IHC slides were photographed under a Nikon Eclipse microscope with a Nikon D7100 camera. All photos were taken under a magnification of 400X for capturing the entire area of hippocampus. Qupath version v2.0 was used for positive stained cell identification and optical density quantification [16]. The background staining threshold RGB values were calibrated consistently on sections treated with the same primary antibody regardless of specimen cohort conditions. The hippocampus was annotated with the polygon wand tool to selectively measure the staining expression within the desired region. Cellular parameters for positive staining detection were fixed based on antibody detection properties. Detected cells was calibrated and quantified by RGB based superpixel clustering. The quantification indicated the average optical density for each detectable cell within the annotated region. An intensity heatmap reflecting the staining quantification was generated for visualization purpose. The gradience of heatmap was adjusted to represent the intensity level across all stained slides that received the same primary antibody.

### Statistics

All numerical data analysis were conducted in Prism (GraphPad Software, La Jolla, CA, USA). 2-way ANOVA was used to compare the population difference between cohorts. P value ≤ 0.05 was considered statistically significant and presented as mean ± SEM. For GSEA, lists of ranked genes based on a score calculated by K-S test as −log10 of P value multiplied by sign of fold-change. Empirical phenotype based 1000 times permutation test produced the nominal P value of the GSEA analysis and prevented the noise from gene-gene correlations.Enrichment score (ES) was calculated by the difference between 0 and the extremes of the entire ranked gene list. The normalized enrichment scores reflected by ES over the average of positive to negative extremes within the gene set [15].

## Results

### The brains from mice treated with a cocktail of Rap, Acb and Pba showed down-regulation of transcriptomic profiles related to each individual drug

The parietal-temporal lobe with hippocampus included from three C57BL/6 mice treated with cocktail and three mice that received control chow were randomly selected for RNA-seq. Total RNA that was purified from homogenized tissue with hippocampus was utilized. All three pathways were suppressed significantly in the brain of mice that received cocktail treatment (Fig.1).

**Figure 1.**
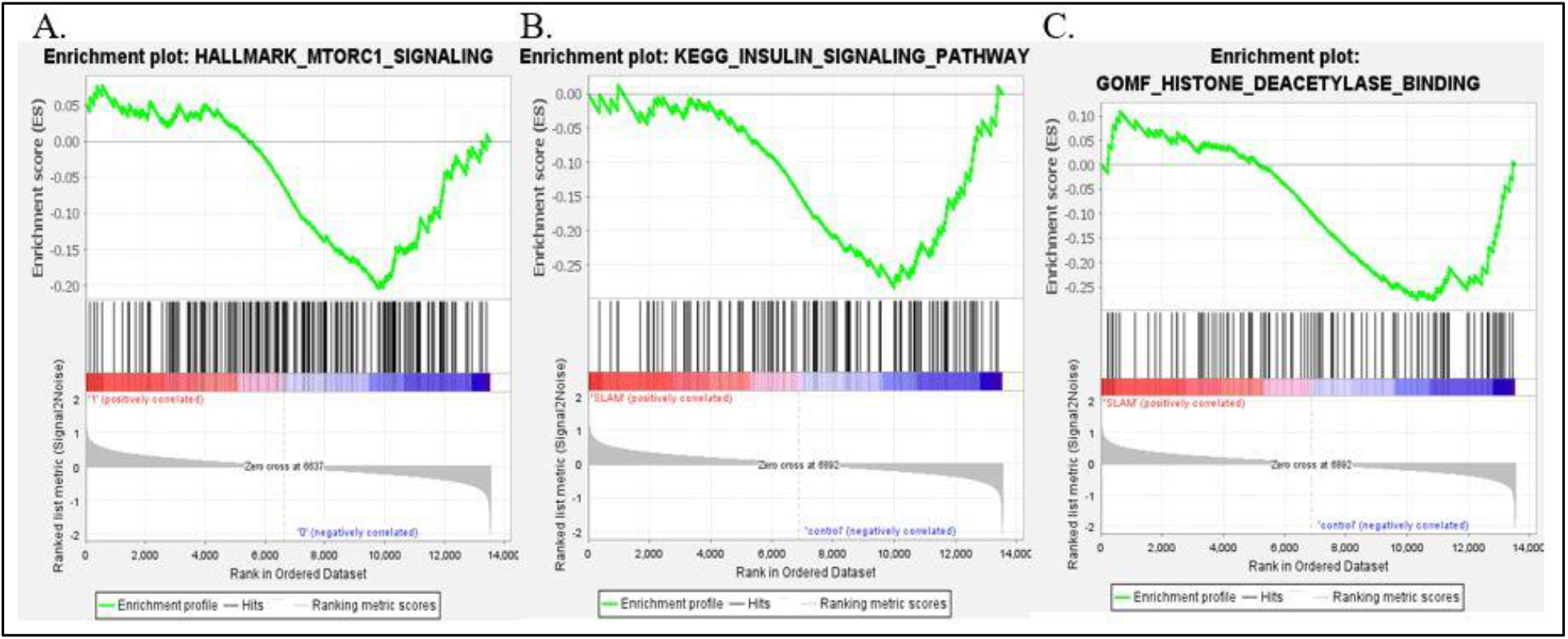
GSEA comparison of brain tissue from cocktail treated and control mice. Results showed that cocktail downregulate mTOR1 signaling, insulin signaling and histone deacetylase binding. A. mTOR1 signaling pathway; B. Insulin signaling pathway; C. Histone deacetylase binding. All nominal p-value ≤ 0.05.

Upon further analysis, a higher false discovery rate q-value (0.38) was observed for the mTOR1 pathway. This could potentially be explained by the nature of the mTOR1 pathway as one of the major nutrient sensing platforms, which is too inclusive and thus contains a large complex of upstream and downstream effectors. The Rap medicated reduction on mTOR1 signaling was reflected by the enhanced expression of downregulating genes. As an anabolic signaling kinase, active mTOR1 translates to rapid cell growth and proliferation. Emerging studies support the idea that lower mTOR1 activity leads to decreased nutrient signaling and extends longevity [17].

The overall insulin signaling referenced to the KEGG database was negatively regulated by the cocktail through enrichment of downregulating or suppressing genes. Insulin signaling is the most conserved aging -controlling pathway in evolution [17-19]. In addition to the beneficiary effect from Rap, the drug Acb promotes glucoregulatory control and improves glucose homeostasis and insulin sensitivity. Current evidence suggests that interventions that constitutively decreased insulin signaling may prevent or reverse the defects associated with aging [20]. Here, induced downstream effectors were seen in the histone deacetylase binding pathway for cocktail treated mice. Mechanistically, Pba targets epigenetic function by inhibiting histone deacetylation, which silences numbers of genes that play a protective role in maintaining systemic as well as neuronal function [21]. Downregulated histone deacetylation alters accessibility of chromatin and allows DNA binding proteins to interact with exposed sites to activate gene transcription and downstream cellular function [22].

### The brains from mice treated with a cocktail of Rap, Acb and Pba showed ameliorated regulation of transcriptomic profiles representing multiple pathways of aging

Brains from mice treated with the cocktail showed a downregulated pattern of DNA repair, suggesting a reduced tendency for DNA damage (Fig. 2A).The DNA repair pathway from GO database include both up and down stream genes involved with DNA repair that makes this an ideal illustration for addressing the level of DNA damage.

**Figure 2.**
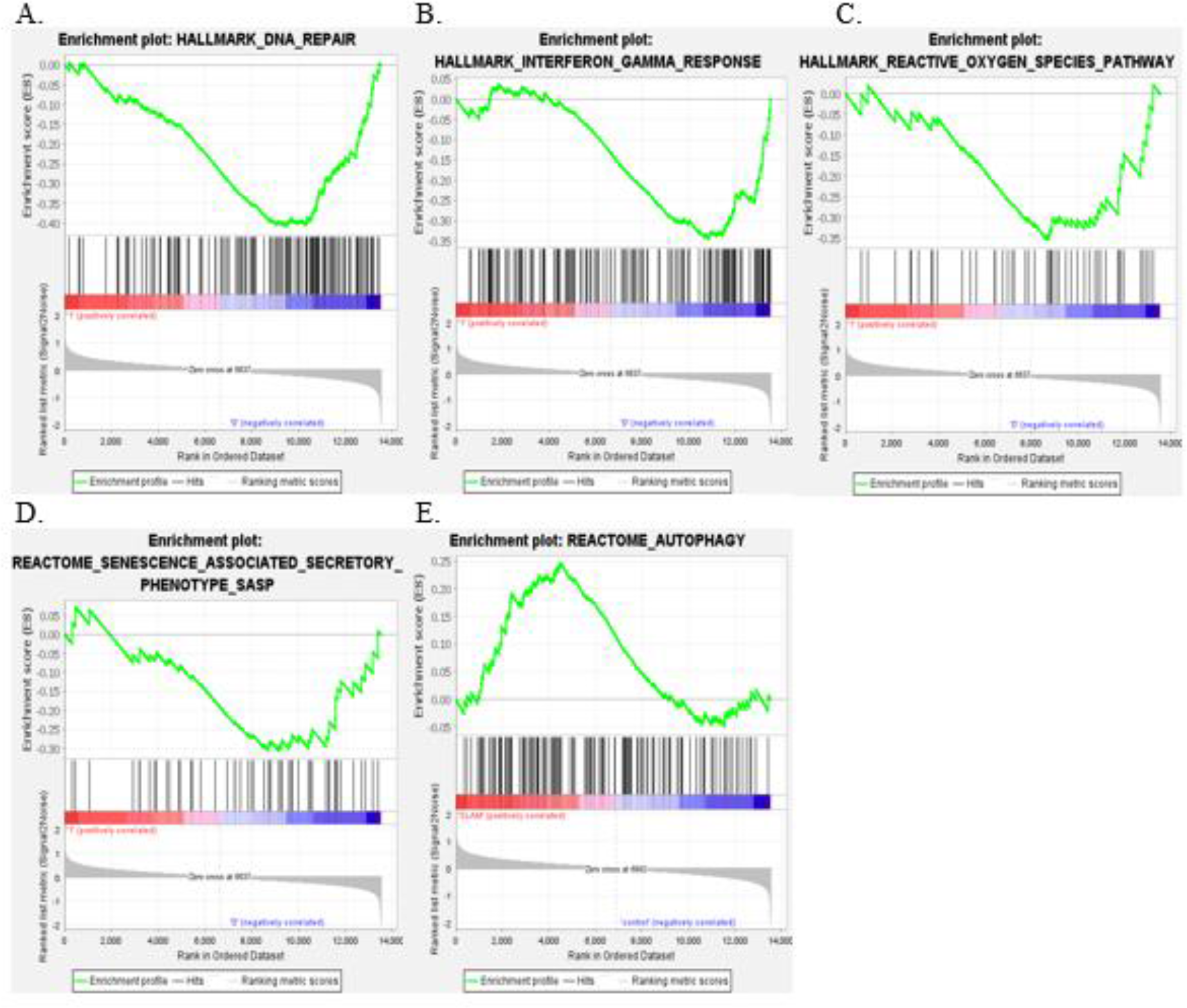
Results for GSEA plots of extended aging pathways. A. All genes involved in DNA repair; B. Genes regulate secretion of interferon-γ; C. Reactive oxygen species (ROS) pathway; D. Genes included for senescence associated secretory phenotype (SASP); E. Genes related to autophagy. All nominal p-value ≤ 0.05 except for SASP (D) and autophagy (E)

Innate immune cytosolic sensing pathway with interferon-γ as downstream effector was chosen to represent primary inflammation response in the CNS [25]. This pathway was down regulated in the brains of mice treated with the cocktail (Fig 2B).

Normal cell division is halted when cellular senescence occurs. As senescence accumulates with increasing age, an elevated expression of both systemic and local senescence associated secretory phenotype (SASP) was observed in neuronal cells [28]. The production of reactive oxygen species (ROS) often promotes cells into senescent phase [29]. ROS and SASP pathways are commonly designated to measure upstream and downstream transcriptional targets of senescence. Both pathways revealed a downregulating trend in the brain of mice treated with the cocktail, however, of these two, only the ROS pathway indicated a statistical significance between the two cohorts (Fig. 2C &D). The GSEA analysis for SASP was referenced to an exclusive Reactome gene database which led to fewer genes presented within the cluster and causing fewer hits from the testing groups. Even with a nominal p-value greater than 0.05, the enrichment plot exhibited a heavily skewed downregulating trend with a distinctive normalized enrichment score of -1.76.

The GSEA results in our study indicated an upregulating tendency for autophagy with data skewed upwards for the mice receiving cocktail diet (Fig. 2D), however, the nominal p-value failed to reject the null. Genomic heatmap interpretation implicated that the K-S ranking for autophagy pathway may be affected by internal variance within treatment cohort.

### Aging pathway protein expression levels corresponded to transcriptomic profiles in brains from mice treated with drug cocktail

Images of IHC stains were processed through Qupath for heatmap demonstration and optical density quantification. The intensity results showed that brains from cocktail treated mice had significantly less expression of γH2AX and HDAC2 while Pba only suppressed the expression of HDAC2 (Fig.3 A&B). HDAC2 is a class I histone deacetylation protein that participates in the DNA damage response as it is rapidly recruited to DNA damage sites to promote hypo-acetylation [31]. γH2AX refers to the X isoform of the histone H2A which resides closely with DNA as a part of the nucleosome. Phosphorylation of γH2AX is an early response to the detection of double-strand breaks or replication stress. The accumulation of HDAC2 and γH2AX in brain tissue have often been inferred to surrogate markers of DNA damage [33].

**Figure 3.**
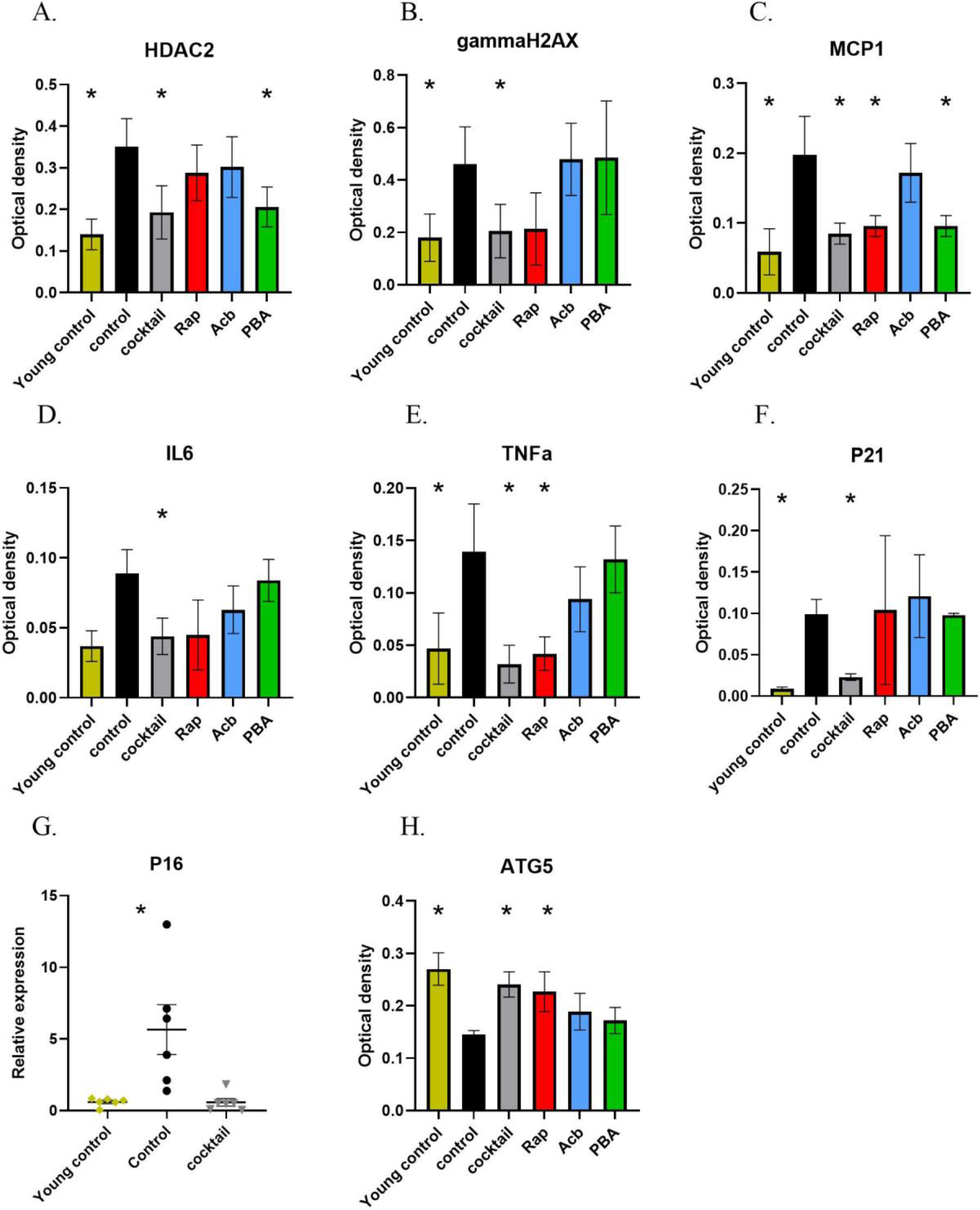
A-G &H: Average optical density value with ± SEM of all positive stained cells in the hippocampus for each tested antibody. * Significant by 2-way ANOVA across each treatment condition, only accepted when P ≤ 0.05. A. HAC2; B. γH2AX; C. MCP-1; D. IL-6; E. TNF-alpha; F. P21; H. ATG5. G: q-PCR value for P16. * Significant accepted as P ≤ 0.05 using 1-way ANOVA.

Reduced levels of all three inflammatory molecules were observed for mice treated with the drug cocktail in our study. Rap alone reduced the level of MCP-1 and TNF-alpha as Pba alone only reduced MCP-1 (Fig.3 C, D&E). General inflammation can be assessed by MCP-1, a pro-inflammatory cytokine that selectively recruits glial cells in the CNS [34]. IL-6 is secreted by parenchymal macrophages in the CNS and is critical for the progression of inflammatory disorders of neural tissue [35]. TNF-alpha is a pro-adhesive inflammatory factor that upregulates NF-κB. Over activation of TNF-alpha induces both acute and chronic neuroinflammation [25].

Biomarkers for senescence were decreased in the brain of cocktail treated mice, and no effect was observed in other treatment cohorts (Fig.3 F&G). P16 and P21 are both well-known endpoints for marking senescence phenotypes in murine models [34]. P21 is a downstream effector of P53 transcription factor and acts as a primary mediator for downstream cell cycle arrest. P16 is essential for oncogene-induced senescence and a cyclin-dependent kinase inhibitor that slows cell division by preventing the progression of the cell cycle from the G1 phase to the S phase [1]. P16 was measured through q-PCR due to lack of validated commercially available antibody for IHC.

ATG5 is a common biomarker for measuring autophagy, which binds with beclin and promotes macroautophagy induction [36]. ATG5 levels were decreased in the brain tissue of both cocktail and Rap treated mice (Fig.3 H).

Heatmaps based on regional optical density provided additional insight on the distribution of staining within the hippocampus region. Staining intensity was visualized by a color gradient, where red represents high levels and blue represents low levels (Fig.4). The heatmap paralleled the quantitative data generated from positive superpixel analysis.

**Figure 4.**
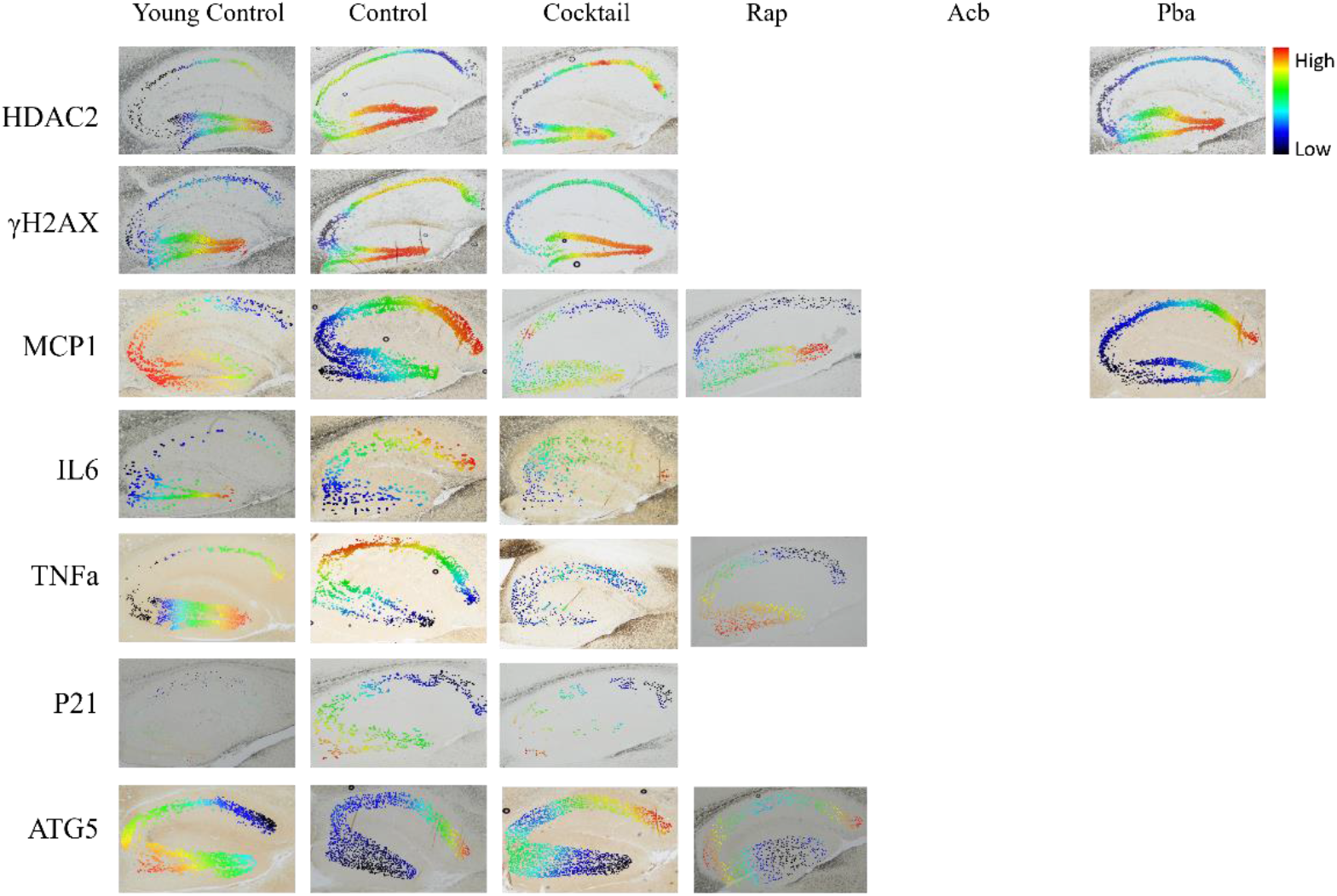
Optical density heatmaps from hippocampal sections imaged at 400X and generated using QuPath digital imaging. Only graphs from cohorts that displayed significant optical density difference compared to control cohort for each antibody are shown. Each row represents staining groups of selected antibodies while each column indicates treatment cohorts. Young controls are listed for negative control demonstration. Brains from young mice are shown as negative controls.

## Discussion

Age related cognitive decline in the brain is characterized by the progressive loss of physiological integrity. This deterioration can lead to early stages of neurodegenerative disease conditions such as Alzheimer’s disease and other dementias, affecting millions of older people [37]. Currently, aging is a subject of scientific scrutiny involving the aggravation of multiple different but interconnected biological processes. Targeting multiple factors that contribute to the progression of aging will likely provide a more robust attenuation of age-related cognitive dysfunction than any single drug [2]. Significant improvement of cognitive performance was observed in middle-aged mice treated for three months with a drug cocktail composed of Rap, Acb and Pba [13]. Selection of these specific drugs for a multiplex approach was based on their well-established roles in promoting health-span and prolonging lifespan in mice [6,9,10].Phenotypic improvement observed from this pre-clinical study suggested the drug cocktail may be targeting multiple pathways associated with aging.

The comparison of mechanistic pathways between the brain tissue of mice receiving drug cocktail treatment versus control were evaluated through GSEA on RNA-seq data. The results enumerated that the drug combination alleviated the aberrant progression of age associated pathways not only directly targeted by each drug but also extensively relevant to overall cocktail effects. Molecular endpoints that manifested among tested aging pathways were further analyzed between all treatment cohorts. Quantification of IHC digital imaging provided additional evidence that the cocktail treatment showed a more comprehensive effect on ameliorating common aspects of aging compared to the administration of each individual drug in the cocktail.

A set of pathways has been considered to characterize the process of aging. Several of these pathways are involved in dysregulated nutrient sensing, enhanced insulin signaling intensity and epigenetic alternations. Activation of mTOR1 signaling, insulin signaling, and histone deacetylase binding pathways can easily antagonize these three biological processes. Emerging studies suggests that Rap, acb and Pba can serve as direct inhibitors for these pathways in various mammalian models [2]. Our results indicated that all three pathways were downregulated in the brain of cocktail treated mice at the transcriptomic level. This finding provides evidence that the orally delivered cocktail was absorbed systemically and effectively into the brain.

Furthermore, the cocktail treatment suppressed the aggravation of many other common aging hallmarks including DNA damage, inflammation and senescence. DNA damage caused by changes to chromatin organization can be mediated through increasing acetylation of histones, suggesting Pba may decrease DNA double-strand breaks or replication stress [23]. mTOR implicates genome integrity maintenance and it has been demonstrated that active mTOR1 expression interplays with DNA damage response systems and promotes growth and survival for cells undergoing constitutive DNA damage [24]. This mechanism aligned with our observation that mice with suppressed mTOR1 signaling and histone deacetylase binding also showed decreased DNA damage level.

mTOR1 is one of the key regulators for chronic inflammatory response as it controls inflammation-associated cell proliferation and migration. Rap as an anti-proliferative mTOR1 inhibitor was proven to prevent endothelial cell overgrowth and is considered a potential reagent for suppressing the primary immune response [26]. Acb was observed to lower glucose excursions and interferes with neuro-inflammatory reactions [27]. This provides the rationale for both Rap and Acb contributing to a lower level of inflammation.

Rap was found to suppress the downstream complexes of mTOR1 signaling including NF-κB, STAT3 and Akt signaling, hence reducing cellular senescence. Increasing evidence implicates that cellular senescence in tissue is associated with dysregulated insulin regulation [30], which suggests that Acb may contribute to reducing the SASP. Moreover, DNA damage is parallel to the production of senescence-associated inflammatory cytokines, which hints to a potential senescence suppressing effect from the administration of Pba. [31].

Autophagy, one of the main activities of the principal proteolytic system implicated in protein quality control, declines with aging. Rap can restore the autophagic flux and enhance autophagy by restoring mitochondrial and lysosomal functions [32]. The phenocopy of caloric restriction from Rap and antihyperglycemic effect from Acb are also known to increase autophagy activation [1]. Interestingly, a trend of upregulated autophagy was observed in the brain of mice that received cocktail but was not statistically significant. This could be explained by the small sample size that increased internal variability within each treatment group.

IHC assessment of age-related molecular endpoints can provide useful information as to how drug treatment can effectively target the expression of specific biomarkers within the brain. Data from this study showed a consistently significant alleviation of expression levels in HDAC2, γH2AX, MCP-1, IL-6, TNF-alpha, P21 and ATG5 from mice treated with cocktail compared to expression levels in control mice. IHC results provided insights of how each of the individual drugs effected the expression of molecular endpoints. Specifically, Rap was almost as effective as the cocktail in ameliorating MCP-1, TNF-alpha and ATG5 levels, while Pba was shown to only decrease HDAC2 and MCP-1 levels. Interestingly, Acb was not effective in attenuating the expression of any selected endpoints but may have contributed to the overall effectiveness of the cocktail. These observations suggest that the drug combination has a major advantage over any of the individual drugs.

All three drugs involved in the cocktail are FDA proved for clinical conditions in people and considered safe with little or no adverse effects. A geropathology assessment in mice from a previous preclinical study using this drug combination showing that the cocktail treatment decreased age-related lesions in all major organs without showing any histology finding related to drug toxicity or abnormal clinical pathology [13].

In conclusion, the framework demonstrated in our study of mapping the alleviating effects of a cocktail of Rap, Acb and Pba applied to aging associated pathways and molecular endpoints provided a cross sectional perspective of mechanistic changes in the brains of old mice. The encouraging observations suggested the cocktail would be a potential therapeutic candidate for preventing, or at the very least, delaying age related cognitive impairment in additional preclinical studies and subsequent clinical trials

## Acknowledgements

Supported in part by NIH grants R01 AG057381 (Ladiges, PI) and R56 AG058543 (Ladiges PI).

## Notes

### Competing Interest Statement

The authors have declared no competing interest.

